# Understanding inter-individual social networks in mixed-species bird flocks

**DOI:** 10.1101/2023.04.13.536662

**Authors:** Akshay Bharadwaj, Aiti Thapa, Akshiti Bhat, Aman Biswakarma, Bharath Tamang, Binod Munda, Biren Biswakarma, Dambar K Pradhan, Dema Tamang, Kabir Pradhan, Mangal K Rai, Pawan Chamling Rai, Rohit Rai, Shambu Rai, Umesh Srinivasan

**Affiliations:** Centre for Ecological Sciences, Indian Institute of Science, Bangalore - 560012, India; Indian Institute of Science Education and Research Kolkata, Mohanpur, Nadia - 741246, West Bengal, India

## Abstract

Mixed-species flocks (MSFs) are an important form of social organisation in forest bird communities worldwide. MSFs provide participants with the benefits of reduced predation risk and/or enhanced foraging efficiency. Recent work has shown that participation in MSFs confers long-term survival benefits in the face of anthropogenic change. However, our understanding of MSFs mainly comes from studies that examine species-level networks, where each node is a unique species and the edges or connections between nodes are associations/interactions between species. While valuable, such approaches might not allow us to understand and investigate the mechanisms that drive MSF formation and structure because social interactions and their effects occur at the individual-level. Empirical studies on multi-species, individual-level MSF social networks have seldom been undertaken due to the various complexities and logistical challenges involved. In this study, we use mist-netting and colour-ringing followed by a standardised observation protocol to construct individual-level social networks in MSFs at 2000m ASL in Eaglenest Wildlife Sanctuary, Arunachal Pradesh, Eastern Himalaya, India. First, we found two separate flocktypes at our study site, comprising two distinct sets of understorey species. The mechanisms contributing to individual-level co-occurrences are likely to differ between these flocktypes, and with MSFs in the Neotropics. The Rusty-fronted Barwing (the nuclear species of one flocktype) shows spatially disjunct territories for each flock while the Yellow-throated Fulvetta (nuclear species of the other flocktype) shows large spatial overlap in its MSF networks, which is likely driven by non-individual-specific benefits such as predation risk dilution. Further, the addition of associating individuals to the social networks has opposite impacts on the two networks. The addition of Coral-billed Scimitar Babblers to the barwing networks greatly reduces network modularity because the associating individuals bridge two modules of barwings (both spatially and in the social network). On the contrary, territorial Rufous-capped Babblers and Grey-cheeked Warblers increase the modularity of the spatially overlapping fulvetta network. Our study provides novel insights into flock formation mechanisms in the Eastern Himalaya, likely applicable to other multi-species flock systems in the Old World.

## Introduction

A society is defined as a cohesive group of individuals that have structured and consistent relationships with each other, which in turn have impacts on their survival and fitness (Farine et al., 2012). A social group is a subunit of a society that is distinguishable from a random aggregation by the persistent and non-random preference or avoidance between individuals (Whitehead, 2008). Direct benefits of intraspecific interactions (like reduction in predation risk) can be hard to distinguish from kin-specific, indirect benefits (like social learning; Clutton-Brock, 2002). The primary mechanisms for the association of non-kin individuals are manipulation and mutualism (Clutton-Brock, 2009). In manipulative interactions, the behaviour and costs for both interacting parties often differ substantially (i.e., there is no change in the beneficiary of this relationship over time). For example, dominant individuals in rhesus macaque societies punish individuals that do not share information when they discover a feeding site (Hauser, 1992). On the other hand, mutualisms are interactions where the interacting individuals receive immediate shared benefits at no overall cost to themselves. For example, in African wild dogs, the cooperation between hunting partners increases the per capita success in catching food (Creel & Creel, 2002).

Social interactions can occur between individuals of the same species or of different species. In the case of interspecific interactions, we can *a priori* exclude kin-based benefits and examine how the direct benefits of social interactions shape social structures and behaviours. Mixed-species bird flocks (hereafter, MSFs) are largely mutualistic foraging associations that consist of individuals from at least two species (Sridhar et al., 2009), and form an important form of social organisation in forest bird communities worldwide (Goodale & Beauchamp, 2010; Thiollay, 1999). MSFs vary greatly in their size, permanence and the strength of associations between species (Moynihan, 1962; Terborgh, 1990). Importantly, mixed-species flocking behaviour has a positive impact on the survival and fitness of participants (Jullien & Clobert, 2000). Further, species that participate in MSFs have been shown to have increased resilience to population declines under long-term anthropogenic change, making this social behaviour relevant for contemporary conservation (Srinivasan, 2019).

Two main mechanisms have been proposed to explain participation in MSFs: reduced predation risk with increasing group sizes and/or improved foraging efficiency. Reduced predation risk can occur through early warning by “sentinel” species (Goodale & Kotagama, 2005; Martínez & Gomez, 2013), the dilution effect (i.e., the reduced probability that a particular individual is depredated on with increasing group size; Foster & Treherne, 1981; Wrona, 1991), the confusion effect (i.e., the reduced ability of a predator to single out an individual from the group; Murali et al., 2019) or the physical disturbance of a predator by flocking birds. Enhanced foraging efficiency can occur through feeding on prey flushed out by other flock participants, learning new foraging locations and avoiding wasteful foraging efforts in patches already exploited by other flock members (Beauchamp, 2005). Although these benefits need not be mutually exclusive (Greenberg, 2001), they can have differential strengths in determining species-specific flocking propensities.

Despite a vast literature on MSF structure and function (Bangal et al., 2022; Harrison & Whitehouse, 2011), our understanding of these systems mainly stems from studying flocks in terms of their interspecific interactions. In other words, our understanding of the social networks in MSFs is largely at the species-level (i.e., each node on the MSF social network corresponds to a flocking species and not an individual; but see Farine & Milburn, 2013). Empirical studies on multi-species, individual-level MSF social networks have seldom been undertaken due to the multiple complexities involved (for instance, the inability to distinguish two individuals of most bird species without employing colour-ringing, necessitating their capture).

Studying MSFs at the individual-level can provide us with a mechanistic understanding of the phenomenon because these interactions and their benefits/costs occur at the level of the individual (Farine et al., 2012). Cooperation theory suggests that heterospecific interactions must occur between non-random individuals to ensure maximised benefits (Clutton-Brock, 2009). Indeed, behavioural studies have shown the presence of heterospecific, individual-level recognition between individuals sharing the same environment (Wascher et al., 2012). Here, we use mist-netting and colour-ringing followed by consistent observations of MSFs to construct detailed individual-level social networks among Eastern Himalayan understorey birds. Building on previous work on species-level associations from the same study area (Borah et al., 2018; Srinivasan, 2019), we asked whether mixed-species flocks comprise individuals that associate consistently with each other across space and time.

## Methods

### Study area

We conducted this study in the montane broadleaf wet evergreen forest at 2000m ASL in the Eaglenest Wildlife Sanctuary, West Kameng district, Arunachal Pradesh, India (27.07°N; 92.40°E). This forest is dominated by *Quercus lamellosa*, *Alnus nipalensis* and *Michelia doltsopa* in the canopy, and bamboo (*Chimonobambusa* sp.) in the understorey (Srinivasan, 2019).

### Mist netting

We conducted mist-netting in four long-term study plots (Fig 1; Srinivasan, 2013). In each plot, a team of 3-4 members operated 20 mist nets (12m length, 4 shelves, 16mm mesh size) from 0630 to 1400h. we systematically set up nets within a plot, with neighbouring nets placed ∼40m apart. We weighed all captured birds, and to enable individual identification without recapture, we ringed each bird with a uniquely numbered aluminium ring and a unique individual-specific colour code. The colour codes consisted of two colour and three colour combinations on specific legs (defined relative to the presence of the aluminium ring; Fig 2A). We conducted mist-netting continuously throughout November and December 2023, except on days when elephants used the study plots. We sampled each plot for three consecutive days before shifting to the next plot. In total, we conducted three rounds of mist-netting in each plot.

**Figure 1:**
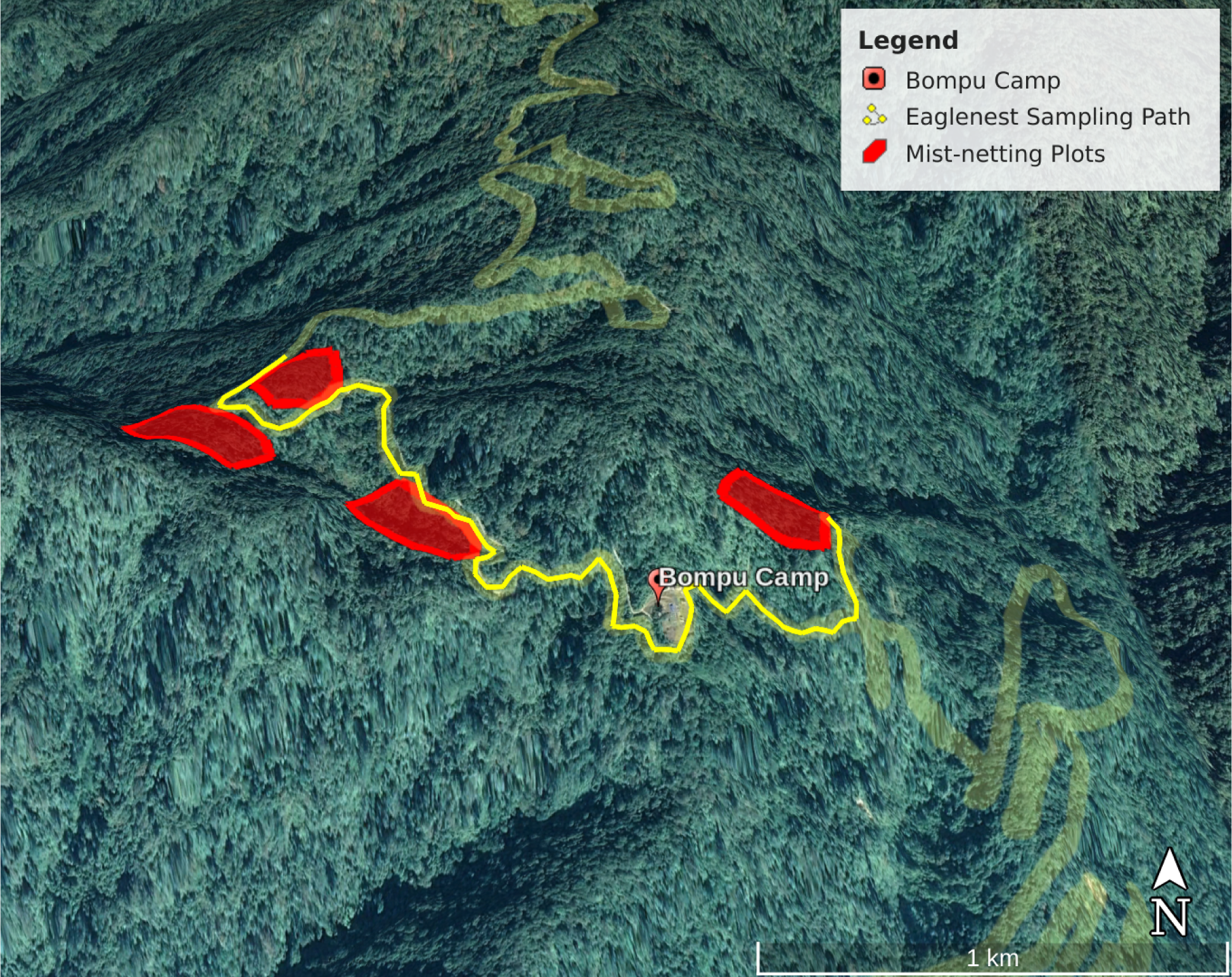
A map representing the mist-netting plots and the stretch of road (bright yellow) that was sampled for MSFs between January and March 2023. Red polygons represent the long-term study plots where we conducted mist-netting.

**Figure 2:**
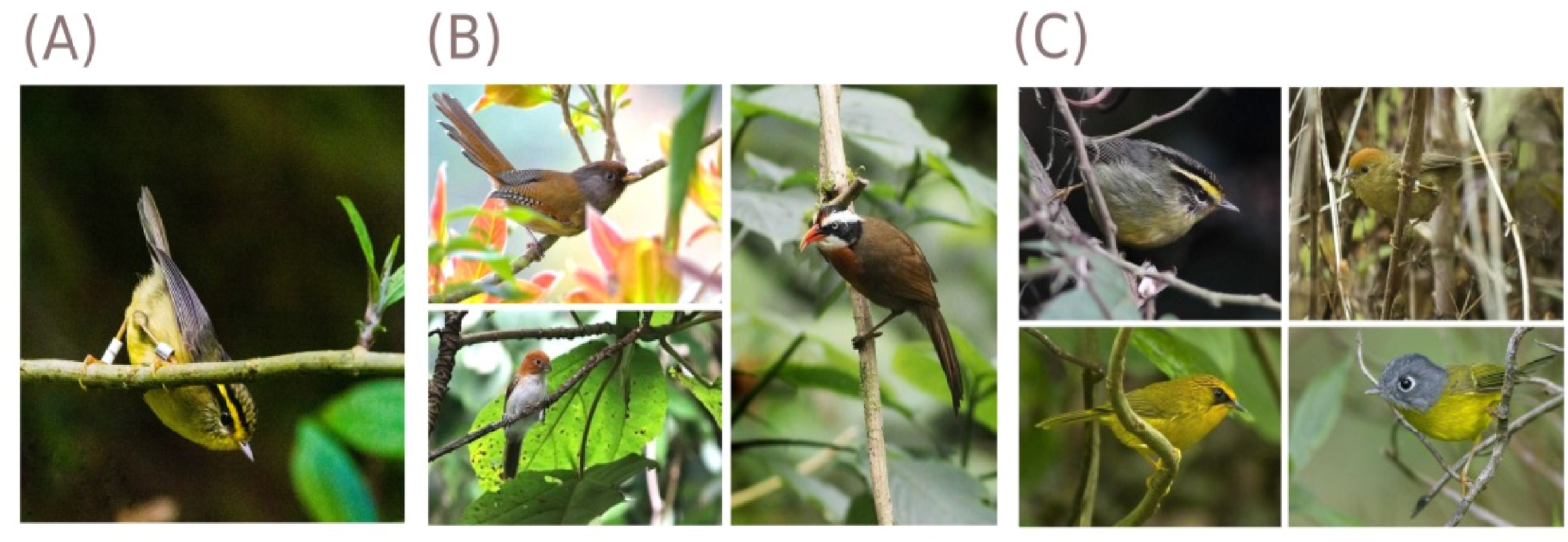
(A) A Yellow-throated Fulvetta (*Schoeniparus cinereus*) colour-ringed with a unique code (White-White) on one leg and has a metal ring on the other leg; (B) Species found in RFBW flocktype are (*clockwise*) Rusty-fronted Barwing (CC-BY-4.0 by Dibyendu Ash), Coral-billed Scimitar Babbler (CC-BY-4.0 by Dibyendu Ash) and Greater Rufous-headed Parrotbill (CC-BY-4.0 by Dibyendu Ash); (C) Species found in YTFV flocktype (*clockwise*) Yellow-throated Fulvetta (CC-BY-4.0 by Dibyendu Ash), Rufous-capped Babbler (CC-BY-SA-2.0 by Francesco Veronesi), Golden Babbler (CC-BY-SA-4.0 by Garima Bhatia) and Grey-cheeked Warbler (CC-BY-SA-3.0 by Dibyendu Ash)

### Mixed-species flock observation

We collected field data on individual co-occurrences between January and March 2023 along a stretch of the fairweather road that passes through Eaglenest Wildlife Sanctuary (Fig 1). For each MSF that we encountered, we noted the colour codes of individuals present in the flock, the species-level composition of the flock, the time of observation and the location of the flock. We collected the location coordinates for each flock using a GARMIN Etrex 20 handheld GPS device. All observations were recorded in a voice recorder (Sony ICD-PX240 MP3 recorder) and only colour codes that we were certain about were noted in the recorder. Note that the species-level flock data consisted of all species seen in the flock, including individuals that were not colour-ringed.

Our initial plan to sample within the plots themselves was relatively unproductive in comparison to sampling along the road because of the speed at which understorey MSFs moved through the undergrowth within the plots. We were unable to follow MSFs for sufficiently long periods of time to reliably record colour codes and species observed in the MSFs. Further, we observed most flocks along the sides of the road, enabling easier and more accurate identification of individual birds through their unique colour ring combinations (example shown in Fig 2A). We conducted consistent sampling along a stretch of the road determined *a priori* (shown in Fig 1).

### Social network analysis (SNA)

#### Species-level social network analysis

We examined social networks at the species-level to understand the various types of flocks (clusters within the species-level networks) that occurred in the study area during winter. To do this, we used the *igraph* package (Csardi & Nepusz, 2006) in the R programming language (R. Development Core Team, 2013) to construct networks at the species-level, and then used a hierarchical clustering algorithm from the package *cluster* (Maechler et al., 2012) to differentiate all the observed flocks into three distinct *flocktypes*. In these networks, each node is a species and edges between nodes are co-occurences between species. Within each flocktype, we defined the nuclear species to be the species with the highest degree (i.e., the most connected with all other species in the flocktype). We used the *dendextend* (Galili, 2015) and *ggplot2* packages (Wickham, 2011) for the cluster dendrogram visualisation.

#### Individual-level social network analysis

First, we filtered the flock dataset to remove any individuals that were potentially misidentified because of difficulty in reading colour codes in the field. Next, we filtered out any flocks which consisted of individuals of only one associating species (i.e., monospecific flocks), although we retained monospecific flocks of nuclear species. Lastly, we removed any individuals that had been recorded less than three times to avoid the effects of individuals with very low-sighting numbers and consequently, low confidence in their social behaviour estimation (Farine & Whitehead, 2015).

For each flocktype, we used the packages *igraph* and *tidyverse* (Wickham et al., 2019) to examine intraspecific and interspecific social networks at the individual-level. In these networks, each node is an individual and edges between nodes are associations between individuals. Using social network analyses, we estimated various network metrics for intra-species (i.e., using data of only the nuclear species) and multi-species flocks (i.e., using data of the nuclear species along with other species present in its flocktype).

#### Spatial distribution of flocks

To understand and visualise the spatial distribution of various subunits (or modules) of the individual-level social networks, we used the *mapview* (Appelhans et al., 2016) and *leaflet* (Cheng et al., 2023) packages in R, and the QGIS software (QGIS.org, 2023). The home range of these modules and their overlap (where necessary) was calculated using the *adeHabitatHR* package (Calenge, 2006).

## Results

### Preliminary results

In total, we recorded 1,007 captures during ∼310 hours of mist-netting across all four study plots. Specifically, we recorded 687 captures of individuals of species that participate in MSFs which we uniquely colour-ringed between November and December 2023.

Subsequently, we observed 317 distinct flocks during our MSF observations between January and March 2023. In total, we recorded co-occurences between 488 individual birds belonging to 18 species. However, only 103 individuals were detected sufficiently frequently to enable their social network analysis (i.e., they passed the different filter thresholds).

### Species-level analysis

Species-level network cluster analysis of MSFs in our study area showed that there are three main flocktypes at our study site:

- Flocktype One consisted of flocks whose central species is the Rusty-fronted Barwing (*Actinodura egertoni)*. Species belonging to this flocktype are Rusty-fronted Barwing, Coral-billed Scimitar Babbler (*Pomatorhinus ferruginosus*) and Greater Rufous-headed Parrotbill (*Psittiparus ruficeps)*. These flocks are hereafter referred to as the **RFBW flocks** (Fig 2B).
- Flocktype Two consisted of flocks whose central species is the Yellow-throated Fulvetta (*Schoeniparus cinereus*). Species belonging to this flocktype are the Yellow-throated Fulvetta, Rufous-capped Babbler (*Cyanoderma ruficeps*), Golden Babbler (*Cyanoderma chrysaeum*) and Grey-cheeked Warbler (*Phylloscopus poliogenys)*. These flocks are hereafter referred to as **YTFV flocks** (Fig 2C).
- Flocktype Three consisted of canopy flocks without any Yellow-throated Fulvetta or Rusty-fronted Barwing individuals.

To analyse only flocks belonging to select flocktypes in the species-level network (shown in Fig S1), the flocks themselves (not species) were clustered into the three different categories (Fig S2). We only analysed flocks that belong only to YTFV and RFBW flocktypes because we did not capture many canopy birds in our understorey mist nets. Therefore, we lacked sufficient individual-level data for the canopy flocktype.

### RFBW flocktype

#### Monospecific flocks

Rusty-fronted Barwings formed distinct modules within their intraspecific social networks showing a high modularity (Fig 3A; N_intra_ = 23, Modularity = 0.69, Density = 0.17). The degree distribution of the network also showed some barwing individuals having a higher degree than most others with near-average degrees (Fig S5).

**Figure 3:**
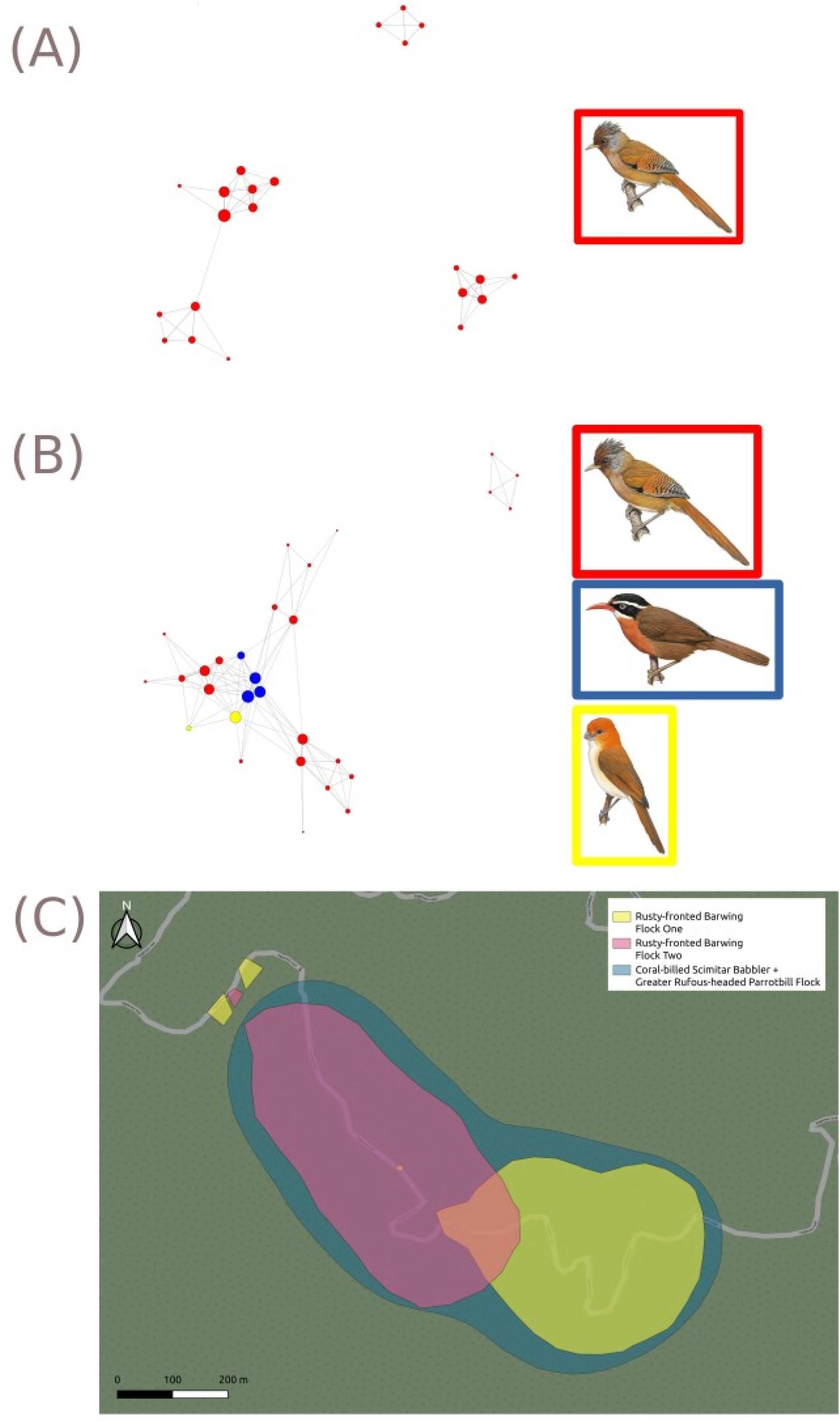
(A) A visualisation of the intraspecies social network between different Rusty-fronted Barwing individuals. Unlike with the Yellow-throated Fulvettas, there are clear modules of associations between distinct individuals. The size of the node shows the degree of the individual in the social network: larger nodes have a larger number of connections with other individuals; (B) A visualisation of the mixed-species social network between different individuals in RFBW flocktype. Red vertices are that of Rusty-fronted Barwing individuals, blue vertices are that of Coral-billed Scimitar Babblers, and yellow vertices are that of Greater Rufous-headed Parrotbills. The size of the node shows the degree of the individual in the social network; larger nodes have a larger number of connections with other individuals; (C) A spatial representation of the “home ranges” of Rusty-fronted Barwing intraspecies units (shown in pink and yellow) with a conserved flock of Greater Rufous-headed Parrotbill and Coral-billed Scimitar Babblers that spans both barwing flock ranges (shown in blue). Note that the “home ranges” are really the region of the road within which the individual or flock-module has been observed.

#### Mixed-species flocks

When other associating species of the RFBW flocktype are added to the network, the modularity of the network is greatly reduced (Fig 3B; N_inter_ = 29, Modularity = 0.39, Density = 0.23). Spatially, we see that the distribution of Barwing flock-modules is largely disjunct from each other, but the range of the Coral-billed Scimitar Babbler flock spans the length of both Rusty-fronted Barwing modules (Fig 3C).

### YTFV flocktype

#### Monospecific flocks of nuclear species

There were two main modules formed in our social network of Yellow-throated Fulvetta individuals. These modules had a high degree of interconnectivity (Fig 4A; N_intra_ = 50, Modularity_intra_ = 0.21, Density_intra_ = 0.24). Some individuals in the network had significantly higher degrees, in comparison to most other individuals with low degrees (Figure S4).

**Figure 4:**
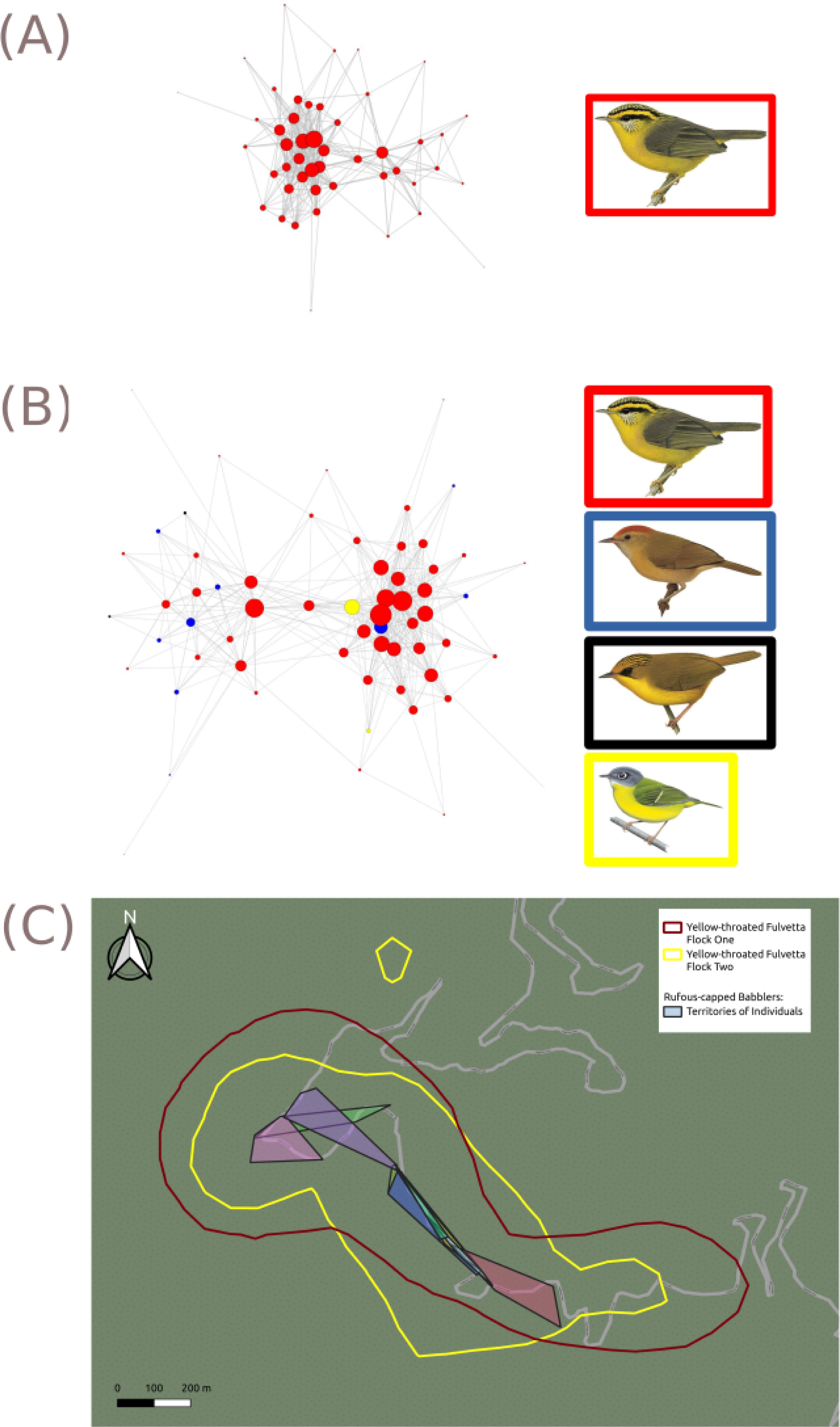
(A) A visualisation of the social network between different Yellow-throated Fulvetta individuals. The size of the node shows the degree of the individual in the social network: larger nodes have a larger number of connections with other individuals; (B) A visualisation of the mixed-species social network between different individuals in YTFV flocktype. Red vertices are that of Yellow-throated Fulvetta individuals, blue vertices are that of Rufous-capped Babblers, yellow vertices are that of Grey-cheeked Warblers, and black vertices are that of Golden Babblers. The size of the node shows the degree of the individual in the social network: larger nodes have a larger number of connections with other individuals; (C) A spatial representation of the “home ranges” of both Yellow-throated Fulvetta modules (shown in yellow and dark red polygons) and different individuals of Rufous-capped Babbler (shaded polygons). Note that the “home ranges” are really the region of the road within which the individual or flock-module has been observed.

#### Mixed-species flocks

When the associating species of the YTFV Flocktype are added into the analyses, we observe that the modularity of the overall network is increased (Fig 4B; N_inter_= 65, Modularity = 0.30, Density = 0.20). Spatially, we found that the modules of Yellow-throated Fulvettas show a high degree of overlap, while Rufous-capped Babblers individuals have fixed territories (Fig 4C).

## Discussion

We show that the flocktypes of understorey birds in the Eastern Himalaya show differential inter-individual association patterns. Our results indicate that the RFBW flocktype exists within territories of monospecific groups of the nuclear species, (i.e, Rusty-fronted Barwing), with the associating species encompassing a larger territory across multiple barwing flock territories. Meanwhile, Yellow-throated Fulvetta flocks are neither highly-modular nor spatially discrete (i.e., high overlap in space use and potentially low territoriality, at least in my sampling locations).

Further, the addition of associating species has opposite effects on the modularity of the two flock networks. The addition of Coral-billed Scimitar Babblers to the presumably territorial subunits of Rusty-fronted Barwings reduces overall network modularity, because a consistent set of individuals of Coral-billed Scimitar Babblers and Greater Rufous-headed Parrotbills connects two distinct, monospecific and spatially segregated flocks of Rusty-fronted Barwings. On the contrary, the modularity of the YTFV flocks is increased by the presence of associating species, likely because of the high territoriality of associating species such as the Rufous-capped Babbler and Grey-cheeked Warbler.

The combined spatial and social network information can help us understand the process of flock formation more mechanistically. The participation of individuals in flocks can be influenced by how different individuals and species use space (Goodale et al., 2015). Our results show that the potential mechanisms underlying MSF behaviour in the mid-elevation Eastern Himalaya are not only different from that of flocktypes shown in the Neotropics, but also from each other.

If the social network of the MSFs formed distinct, non-interconnected modules that are also spatially disjunct from each other across all the species in the flocktype, then there would have been multi-species territoriality of MSFs as in the Amazon basin. In the Amazonian MSFs, a species-level study observed that mixed-species bird flocks are territorial at the flock level; each MSF is composed of a pair of individuals of each species, which defend a common territory as a single multi-species unit (Martínez & Gomez, 2013; Munn & Terborgh, 1979). Studies at the individual-level have subsequently confirmed this multi-species territorial behaviour, with intraspecific aggression between individuals at the boundaries of two mixed-species flock territories (Jullien & Thiollay, 1998).

If the social networks of the MSFs form distinct, non-interconnected modules that are spatially segregated within each species but not across all the species in a flocktype (i.e., each intraspecies flock/individual defends a territory, but not at the multi-species level), then the driver behind which individual flocks with which other individual is likely to simply be spatially-driven associations between individuals of different species depending on the distributions of their territories. In Panama, for instance, five passerine bird species tended to join flocks whenever one was available inside their home range, regardless of the flock’s specific location within that home range (Pomara et al., 2007). Notably, their five study species were “flock joiners”, or associating species, which possessed territories and joined flocks which had other nuclear species (Pomara et al., 2007). Similar to patterns in Panamanian flocks, Rufous-capped Babbler and Grey-cheeked Warbler are associating species that appear to hold specific territories, and associate with flocks of the nuclear species, Yellow-throated Fulvetta, that pass through their territory. On the contrary, Rusty-fronted Barwings (the nuclear species of the RFBW flocktype) are territorial in nature and consist of monospecific flocks of possibly related individuals (with kin selection likely to be an important driver of benefit sharing between individuals). Meanwhile, the associating species, such as the Coral-billed Scimitar Babbler and the Greater Rufous-headed parrotbills have territories that span a larger area, which encompasses the territory of multiple barwing flocks. Here, the nuclear species is territorial and forms flocks with individuals of associating species passing through their territory.

If the social networks of the MSFs form highly interconnected modules that are spatially overlapping, the drivers behind co-occurrences between individuals are likely not related to kin selection but arise from non-individual-specific mechanisms such as predation risk dilution. This is the pattern we observe in the YTFV flocktype in Eaglenest, where the Yellow-throated Fulvetta is the nuclear species and appears to have random inter-individual associations. This pattern is contradictory to the previously reported non-random associations between individuals (i.e, with a preference for certain individuals over others), despite overlapping home ranges between multiple individuals/flock-modules (Farine & Milburn, 2013).

Yellow-throated fulvetta groups exhibit fission-fusion dynamics, a social structure found in a wide range of species (Kutsukake, 2006; Micheletta et al., 2012). Life-history traits such as communal roosting and colonial nesting among birds drive fission-fusion dynamics at different temporal scales (Silk et al., 2014). Further, fission-fusion dynamics have been reported to occur in situations with spatial variability in environment and resource availability (Sueur et al., 2011). For example, migratory birds such as the Egyptian Vulture do not form aggregations in their wintering grounds (where carcasses are abundant) but do so in Europe, where resources are localised and scarce (Cortés-Avizanda et al., 2011). Environmental variation across a diel cycle can also lead to differences in community structure viz. changes in predator/prey communities over the course of the day (Farine & Lang, 2013; Silk et al., 2014). The resulting changes could be individual-specific (depending on differential body condition) and lead to altered social behaviour, in light of changing foraging-predation tradeoffs. Using the temporally dynamic network approaches (Cantor et al., 2012; Fig S6), we can examine the stability of social structures across various timescales.

The social networks of barwings and fulvettas differ considerably in terms of their structure and modularity. There are potentially two broad reasons behind this difference, relating to the nature of MSF benefits themselves. The first is potentially different dietary resources consumed by the species and the distribution of these resources (Cortés-Avizanda et al., 2011; Johnson et al., 2002). The territorial barwing flocks may have distinct territories and family groups due to a highly-specialised diet of resources that are spatially clustered (Grant, 1993). This could lead to the preference of defending a territory and resource patches with your kin. Along similar lines, we expect fulvettas to rely on spatially distributed resources (chiefly, understorey arthropods) which do not necessitate the costs of defending territories outside of the breeding season (Grant, 1993).

Another potential driver of differential social structures between the two flocktypes could be the role of predator detection and alarm calling (Martínez et al., 2016). Barwing individuals may have assigned roles in detecting predators, with some individuals playing the role of sentinels by signalling alarm calls while others forage. Similar to meerkat societies (Rauber & Manser, 2018), barwings may form groups where the benefits of flock formation are complementary and shared with their kin (Goodale et al., 2020; Griesser & Ekman, 2004). On the other hand, in yellow-throated fulvettas, the individual’s ability to detect their predators could be lower due to their myopic understorey foraging behaviour, resulting in no distinct roles of sentinel and foraging individuals. As a result, fulvettas could prioritise being part of a flock to receive the supplementary benefits of per capita predation risk dilution, without much heed to which individual identities of its members (Goodale et al., 2020).

The effect of kin-specific benefits on determining individual co-occurrences can be investigated further through the estimation of relatedness between individuals. Our results show that in the RFBW flocktype, consistent sets of individuals occupy a fixed territory. Using parentage and relatedness assays based on large numbers of single nucleotide polymorphisms (SNP) loci can help us understand the genetic relationships between many individuals (Thrasher et al., 2018). For instance, in House Sparrows (*Passer domesticus*), individuals prefer to flock with same-brood siblings during social activities in comparison to non-sib birds (Tóth et al., 2009). Individuals/groups of other species could then “eavesdrop” on communications between kin to access resources or avoid predation (Goodale & Kotagama, 2005; Martínez et al., 2022).

When faced with environmental change, most vertebrates respond physiologically by releasing stress hormones such as glucocorticoids (like corticosterone, i.e., CORT). These hormones, in turn, elicit physiological and behavioural changes commonly referred to as the “stress response” (Blas et al., 2007). Estimating CORT levels in slow-developing and highly vascularised feathers provides a measure of chronic stress over longer timeframes (i.e., the growing period of the feather). Estimating the CORT levels of various flocking individuals is an exciting avenue for future research, allowing an examination of correlations between stress, individual roles in networks (degree, centrality), and survival rates over time.

Lastly, from a methodology perspective, the use of RFID-based sensors can help us identify individuals more accurately and reduce the probability of misidentification or non-detection of tagged birds. Several studies on temperate forest birds use a mist-netting protocol that involves attaching Passive Integrated Transponder or PIT tags onto the birds’ tarsi (Farine & Lang, 2013). These tags make them detectable when they are in proximity to loggers with antennae. Importantly, these electronic tags and automated data collection can provide complete or near-complete datasets of individual co-occurrences (Farine & Whitehead, 2015).

Anthropogenic habitat changes likely have a considerable impact on the social systems of birds. Comparative studies of mixed-species flocks in urban and forest habitats in Malaysia show differing social dynamics (Lee et al., 2005). Therefore, coupled with abiotic predictors of species’ vulnerabilities (Bharadwaj et al., 2022), understanding the species-specific drivers of MSF formation (and how these might change with habitat modification) helps us better understand which species are at a greater extinction risk under current anthropogenic changes. To do this, examining MSF behaviour at the individual-level, where co-occurences between conspecific individuals can be distinguished and recorded, can provide greater conservation insights than species-level patterns. Our study provides novel insights into flock formation mechanisms in the Eastern Himalaya, likely applicable to other multi-species flock systems in the Old World.

**Table 1:** A list of all calculated metrics, their levels, definitions and biological significance (Brandes et al., 2008; Lau et al., 2017)

| Metrics used | Level | Definition | Biological significance | Range of values |
| --- | --- | --- | --- | --- |
| Number of Nodes (N) | Network property | Total number of nodes present in the network | Number of individuals present in the social network analysis | $[1, \infty] \in \mathbb{Z}^+$ |
| Modularity | Network property | Degree to which edges are distributed within rather than between distinct sets of nodes | Indicates the degree of presence/absence of distinct sub-units within the social network (distinct flocks) | $[-0.5, 1] \in \mathbb{R}$ |
| Density | Network property | The proportion of possible edges that are actually observed in the network | Quantifies the interconnectedness of all individuals in the network. High network density implies a network where individuals associate with many other individuals. | $[0, 1] \in \mathbb{R}^+$ |
| Degree | Node property | Number of edges connected to a given node, which is a type of local centrality | Indicates how social an individual is. A high degree implies the individual is found to associate with many other individuals. | $[1, N] \in \mathbb{Z}^+$<br>where **N** is the number of nodes in the network |

## Acknowledgements

We also thank the funding organisations that funded our study, the Indian Institute of Science, Science and Engineering Research Board, GoI, Department of Biotechnology, GoI and the Wild Animal Initiative. My thanks also to the Arunachal Pradesh Forest Department for the permits to conduct mark-resight studies in the Eaglenest Wildlife Sanctuary, Arunachal Pradesh.

## Supplementary material

**Figure S1:**
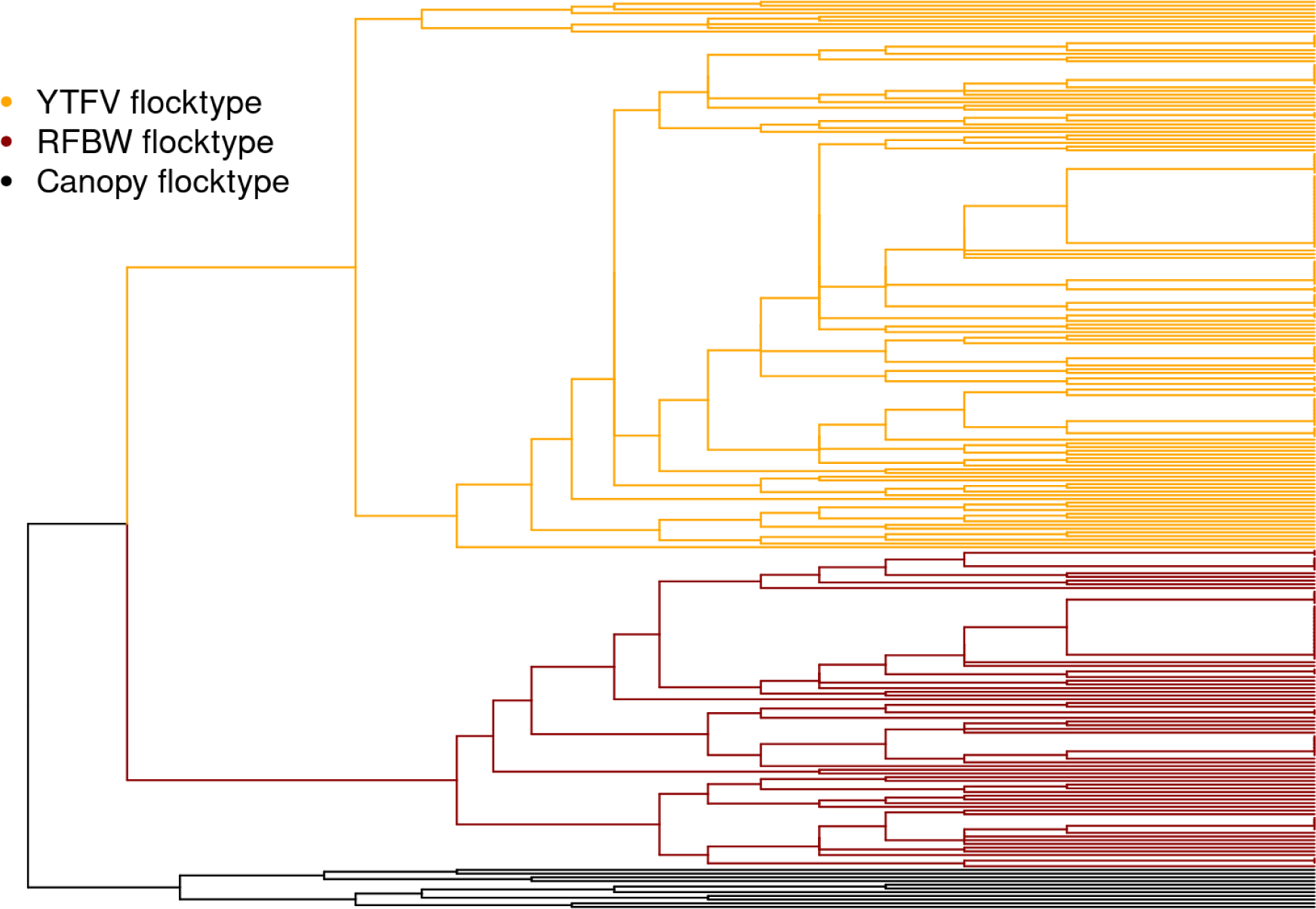
A cluster dendrogram showing the observed flocks clustered according to their similarity with each other, and into three different categories, i.e, flocktypes. YTFV flocktype (yellow) consists of understorey insectivorous birds led by Yellow-throated Fulvetta. RFBW flocktype (dark red) consists of flocks led by Rusty-fronted Barwings. Canopy flocktype (black) consists of canopy flocks led by Yellow-cheeked Tit (*Machlolophus spilonotus*) and Yellow-browed Tit (*Sylviparus modestus*).

**Figure S2:**
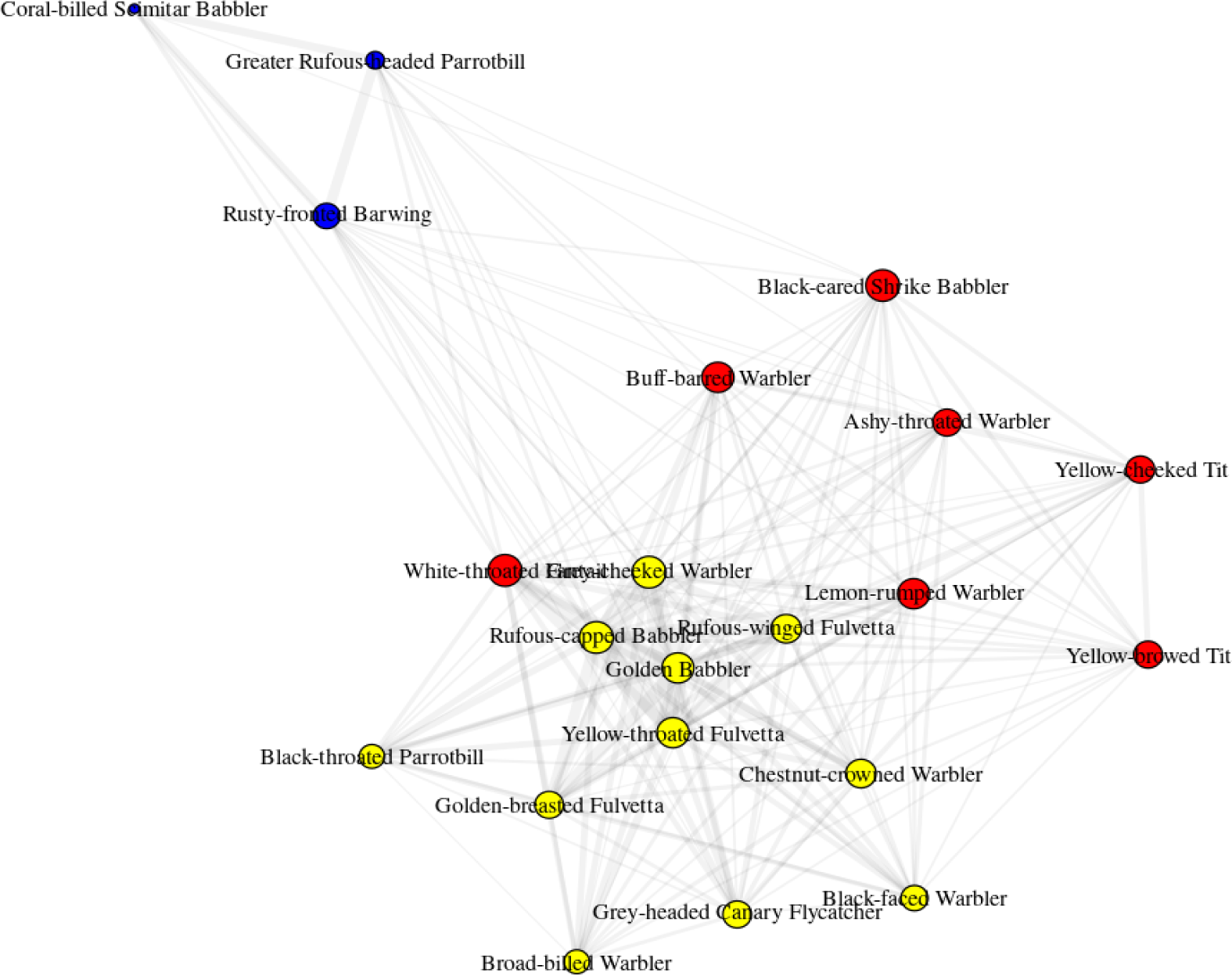
Species-level network showing the presence of three distinct clusters, corresponding to different flocktypes in our dataset.

**Figure S3:**
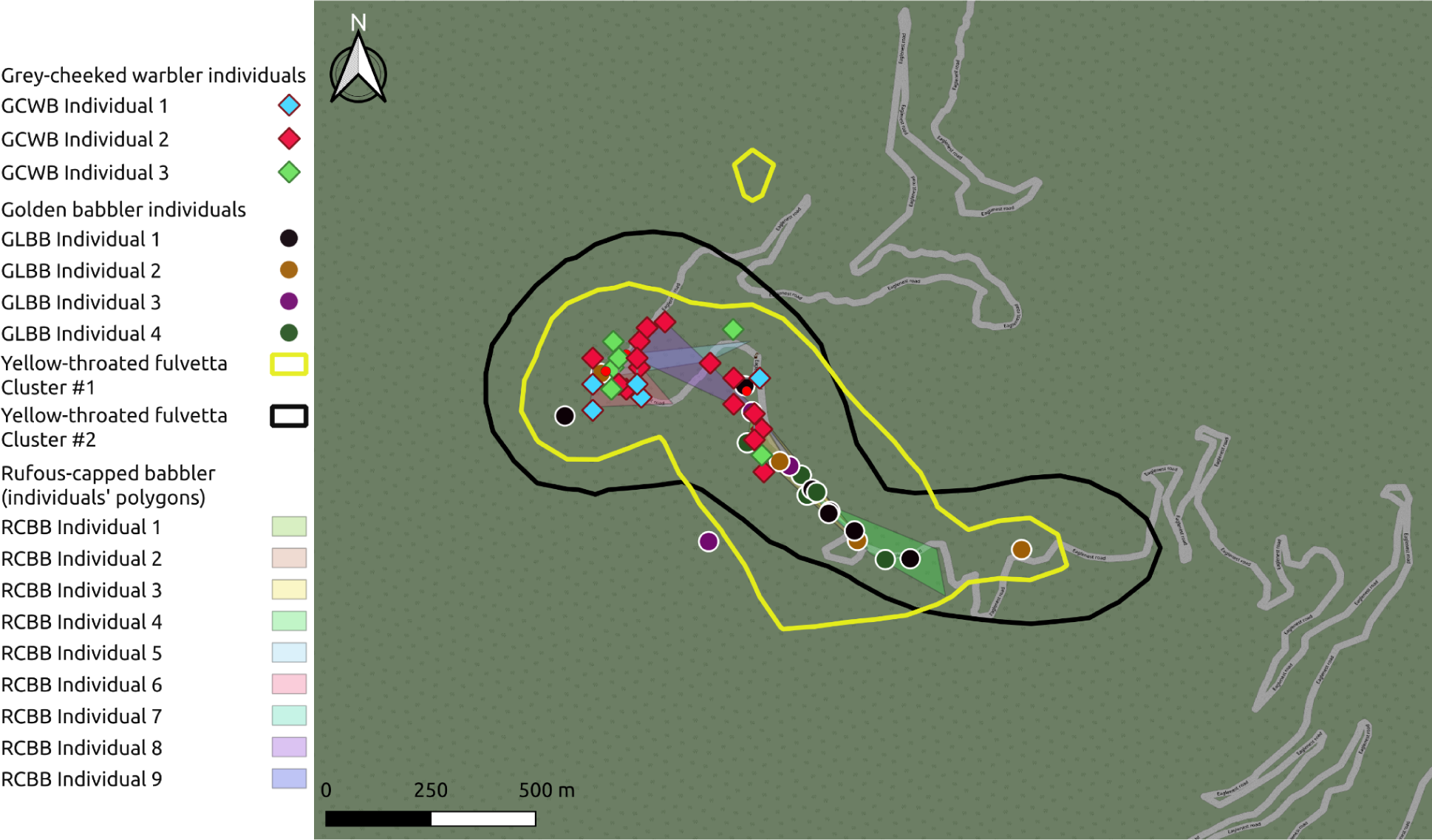
A spatial representation of the “home ranges” of both Yellow-throated Fulvetta modules (shown in yellow and black), the spatial distribution of Grey-cheeked Warbler (diamonds), Rufous-capped Babbler (shaded polygons) and Golden Babbler individuals (filled circles). Note that the “home ranges” are really the region of the road within which the individual or flock-module has been observed.

**Figure S4:**
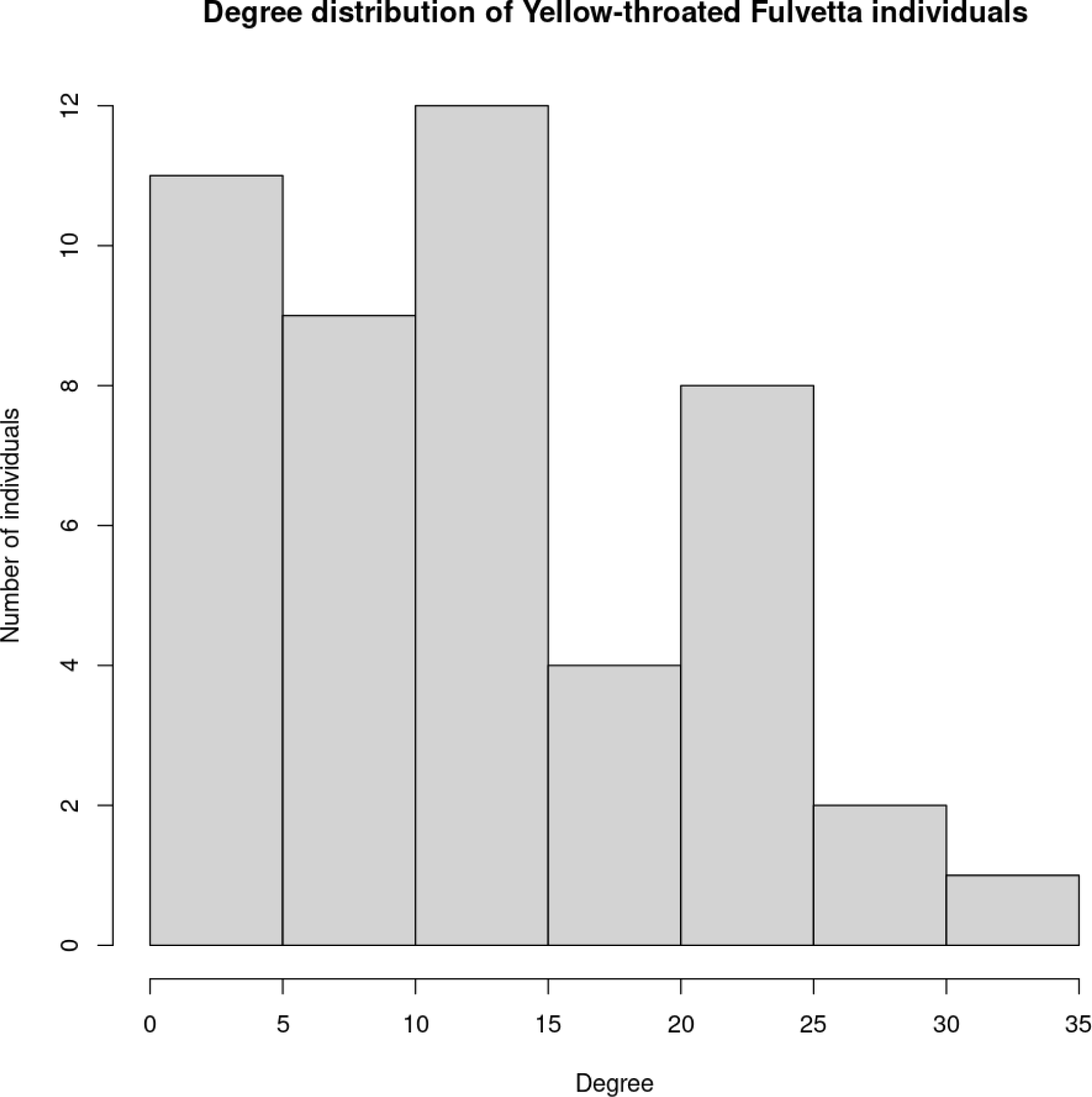
Degree distribution for Yellow-throated Fulvetta individuals.

**Figure S5:**
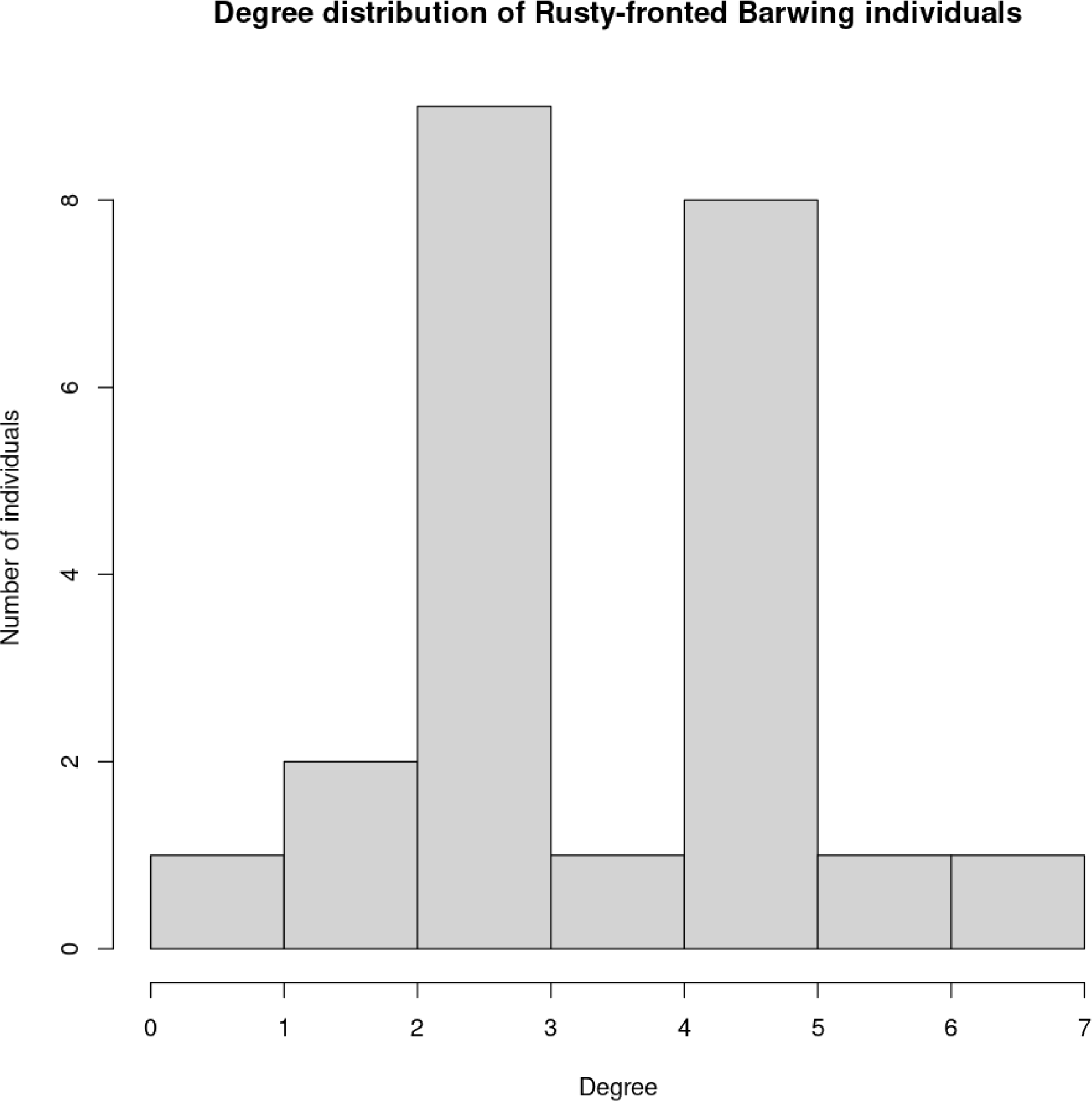
Degree distribution for Rusty-fronted Barwing individuals.

**Figure S6:** A visualisation of interactions in the YTFV flocktype over the course of the field season. https://drive.google.com/file/d/1Ak72ypD1UuyKy2YhRFEUrFbKDzvFyy8a/view?usp=sharing.

## References

Appelhans, T., Detsch, F., Reudenbach, C., & Woellauer, S. (2016). Mapview-Interactive viewing of spatial data in R. EGU General Assembly Conference Abstracts, EPSC2016-1832.

Bangal, P., Sridhar, H., Shizuka, D., Vander Meiden, L. N., & Shanker, K. (2022). Flock-species richness influences node importance and modularity in mixed-species flock networks. Oecologia, 198(2), 431–440. 10.1007/s00442-021-05053-z

Beauchamp, G. (2005). Does group foraging promote efficient exploitation of resources? Oikos, 111(2), 403–407. 10.1111/j.0030-1299.2005.14136.x

Bharadwaj, A., Chanda, R., & Srinivasan, U. (2022). Abiotic niche predictors of long-term trends in body mass and survival of Eastern Himalayan birds (p. 2022.08.25.505219). bioRxiv. 10.1101/2022.08.25.505219

Blas, J., Bortolotti, G. R., Tella, J. L., Baos, R., & Marchant, T. A. (2007). Stress response during development predicts fitness in a wild, long lived vertebrate. Proceedings of the National Academy of Sciences, 104(21), 8880–8884. 10.1073/pnas.0700232104

Borah, B., Quader, S., & Srinivasan, U. (2018). Responses of interspecific associations in mixed-species bird flocks to selective logging. Journal of Applied Ecology, 55(4), 1637– 1646. 10.1111/1365-2664.13097

Brandes, U., Delling, D., Gaertler, M., Gorke, R., Hoefer, M., Nikoloski, Z., & Wagner, D. (2008). On Modularity Clustering. IEEE Transactions on Knowledge and Data Engineering, 20(2), 172–188. 10.1109/TKDE.2007.190689

Calenge, C. (2006). The package “adehabitat” for the R software: A tool for the analysis of space and habitat use by animals. Ecological Modelling, 197(3–4), 516–519.

Cantor, M., Wedekin, L. L., Guimarães, P. R., Daura-Jorge, F. G., Rossi-Santos, M. R., & Simões-Lopes, P. C. (2012). Disentangling social networks from spatiotemporal dynamics: The temporal structure of a dolphin society. Animal Behaviour, 84(3), 641– 651. 10.1016/j.anbehav.2012.06.019

Cheng, J., Karambelkar, B., Xie, Y., Wickham, H., Russell, K., Johnson, K., Schloerke, B., & Agafonkin, V. (2023). Package ‘leaflet.’

Clutton-Brock, T. (2002). Breeding together: Kin selection and mutualism in cooperative vertebrates. Science, 296(5565), 69–72.

Clutton-Brock, T. (2009). Cooperation between non-kin in animal societies. Nature, 462(7269), Article 7269. 10.1038/nature08366

Cortés-Avizanda, A., Almaraz, P., Carrete, M., Sánchez-Zapata, J. A., Delgado, A., Hiraldo, F., & Donázar, J. A. (2011). Spatial Heterogeneity in Resource Distribution Promotes Facultative Sociality in Two Trans-Saharan Migratory Birds. PLOS ONE, 6(6), e21016. 10.1371/journal.pone.0021016

Creel, S., & Creel, N. M. (2002). The African wild dog: Behavior, ecology, and conservation (Vol. 25). Princeton University Press.

Csardi, G., & Nepusz, T. (2006). The igraph software package for complex network research. *InterJournal*, Complex Systems, 1695(5), 1–9.

Farine, D. R., Garroway, C. J., & Sheldon, B. C. (2012). Social network analysis of mixed-species flocks: Exploring the structure and evolution of interspecific social behaviour. Animal Behaviour, 84(5), 1271–1277. 10.1016/j.anbehav.2012.08.008

Farine, D. R., & Lang, S. D. J. (2013). The early bird gets the worm: Foraging strategies of wild songbirds lead to the early discovery of food sources. Biology Letters, 9(6), 20130578. 10.1098/rsbl.2013.0578

Farine, D. R., & Milburn, P. J. (2013). Social organisation of thornbill-dominated mixed-species flocks using social network analysis. Behavioral Ecology and Sociobiology, 67(2), 321–330. 10.1007/s00265-012-1452-y

Farine, D. R., & Whitehead, H. (2015). Constructing, conducting and interpreting animal social network analysis. Journal of Animal Ecology, 84(5), 1144–1163. 10.1111/1365-2656.12418

Foster, W. A., & Treherne, J. E. (1981). Evidence for the dilution effect in the selfish herd from fish predation on a marine insect. Nature, 293(5832), 466–467.

Galili, T. (2015). dendextend: An R package for visualizing, adjusting and comparing trees of hierarchical clustering. Bioinformatics, 31(22), 3718–3720.

Goodale, E., & Beauchamp, G. (2010). The relationship between leadership and gregariousness in mixed-species bird flocks. Journal of Avian Biology, 41(1), 99–103. 10.1111/j.1600-048X.2009.04828.x

Goodale, E., Ding, P., Liu, X., Martínez, A., Si, X., Walters, M., & Robinson, S. K. (2015). The structure of mixed-species bird flocks, and their response to anthropogenic disturbance, with special reference to East Asia. Avian Research, 6(1), 14. 10.1186/s40657-015-0023-0

Goodale, E., & Kotagama, S. W. (2005). Alarm Calling in Sri Lankan Mixed-Species Bird Flocks. The Auk, 122(1), 108–120. 10.1093/auk/122.1.108

Goodale, E., Sridhar, H., Sieving, K. E., Bangal, P., Colorado Z., G. J., Farine, D. R., Heymann, E. W., Jones, H. H., Krams, I., Martínez, A. E., Montaño-Centellas, F., Muñoz, J., Srinivasan, U., Theo, A., & Shanker, K. (2020). Mixed company: A framework for understanding the composition and organization of mixed-species animal groups. Biological Reviews, 95(4), 889–910. 10.1111/brv.12591

Grant, J. W. A. (1993). Whether or not to defend? The influence of resource distribution. Marine Behaviour and Physiology, 23(1–4), 137–153. 10.1080/10236249309378862

Greenberg, R. S. (2001). Birds of many feathers: The formation and structure of mixed species flocks of forest birds. On the Move: How and Why Animals Travel in Groups.

Griesser, M., & Ekman, J. (2004). Nepotistic alarm calling in the Siberian jay, Perisoreus infaustus. Animal Behaviour, 67(5), 933–939. 10.1016/j.anbehav.2003.09.005

Harrison, N. M., & Whitehouse, M. J. (2011). Mixed-species flocks: An example of niche construction? Animal Behaviour, 81(4), 675–682.

Hauser, M. D. (1992). Costs of deception: Cheaters are punished in rhesus monkeys (Macaca mulatta). Proceedings of the National Academy of Sciences, 89(24), 12137–12139.

Johnson, D. D. P., Kays, R., Blackwell, P. G., & Macdonald, D. W. (2002). Does the resource dispersion hypothesis explain group living? Trends in Ecology & Evolution, 17(12), 563–570. 10.1016/S0169-5347(02)02619-8

Jullien, M., & Clobert, J. (2000). The survival value of flocking in Neotropical birds: Reality or fiction? Ecology, 81(12), 3416–3430.

Jullien, M., & Thiollay, J.-M. (1998). Multi-species territoriality and dynamic of neotropical forest understorey bird flocks. Journal of Animal Ecology, 67(2), 227–252. 10.1046/j.1365-2656.1998.00171.x

Kutsukake, N. (2006). The Context and Quality of Social Relationships Affect Vigilance Behaviour in Wild Chimpanzees. Ethology, 112(6), 581–591. 10.1111/j.1439-0310.2006.01200.x

Lau, M. K., Borrett, S. R., Baiser, B., Gotelli, N. J., & Ellison, A. M. (2017). Ecological network metrics: Opportunities for synthesis. Ecosphere, 8(8), e01900. 10.1002/ecs2.1900

Lee, T. M., Soh, M. C. K., Sodhi, N., Koh, L. P., & Lim, S. L.-H. (2005). Effects of habitat disturbance on mixed species bird flocks in a tropical sub-montane rainforest. Biological Conservation, 122(2), 193–204. 10.1016/j.biocon.2004.07.005

Maechler, M., Rousseeuw, P., Struyf, A., Hubert, M., & Hornik, K. (2012). Cluster: Cluster analysis basics and extensions.

Martínez, A. E., & Gomez, J. P. (2013). Are Mixed-Species Bird Flocks Stable through Two Decades? The American Naturalist, 181(3), E53–E59. 10.1086/669152

Martínez, A. E., Gomez, J. P., Ponciano, J. M., & Robinson, S. K. (2016). Functional Traits, Flocking Propensity, and Perceived Predation Risk in an Amazonian Understory Bird Community. The American Naturalist, 187(5), 607–619. 10.1086/685894

Martínez, A. E., Parra, E., Gomez, J. P., & Vredenburg, V. T. (2022). Shared predators between primate groups and mixed species bird flocks: The potential for forest-wide eavesdropping networks. Oikos, 2022(10), e08274. 10.1111/oik.08274

Micheletta, J., Waller, B. M., Panggur, M. R., Neumann, C., Duboscq, J., Agil, M., & Engelhardt, A. (2012). Social bonds affect anti-predator behaviour in a tolerant species of macaque, Macaca nigra. Proceedings of the Royal Society B: Biological Sciences, 279(1744), 4042–4050. 10.1098/rspb.2012.1470

Moynihan, M. H. (1962). The organization and probable evolution of some mixed species flocks of neotropical birds.

Munn, C. A., & Terborgh, J. W. (1979). Multi-Species Territoriality in Neotropical Foraging Flocks. The Condor, 81(4), 338–347. 10.2307/1366956

Murali, G., Kumari, K., & Kodandaramaiah, U. (2019). Dynamic colour change and the confusion effect against predation. Scientific Reports, 9(1), Article 1. 10.1038/s41598-018-36541-7

Pomara, L. Y., Cooper, R. J., & Petit, L. J. (2007). Modeling the flocking propensity of passerine birds in two Neotropical habitats. Oecologia, 153(1), 121–133. 10.1007/s00442-007-0701-7

QGIS.org. (2023). QGIS Geographic Information System [Computer software]. http://www.qgis.org/

R. Development Core Team. (2013). R: A language and environment for statistical computing.

Rauber, R., & Manser, M. B. (2018). Experience of the signaller explains the use of social versus personal information in the context of sentinel behaviour in meerkats. Scientific Reports, 8(1), Article 1. 10.1038/s41598-018-29678-y

Silk, M. J., Croft, D. P., Tregenza, T., & Bearhop, S. (2014). The importance of fission–fusion social group dynamics in birds. Ibis, 156(4), 701–715. 10.1111/ibi.12191

Sridhar, H., Beauchamp, G., & Shanker, K. (2009). Why do birds participate in mixed-species foraging flocks? A large-scale synthesis. Animal Behaviour, 78(2), 337–347. 10.1016/j.anbehav.2009.05.008

Srinivasan, U. (2013). A slippery slope: Logging alters mass–abundance scaling in ecological communities. Journal of Applied Ecology, 50(4), 920–928. 10.1111/1365-2664.12123

Srinivasan, U. (2019). Morphological and Behavioral Correlates of Long-Term Bird Survival in Selectively Logged Forest. Frontiers in Ecology and Evolution, 7. https://www.frontiersin.org/article/10.3389/fevo.2019.00017

Sueur, C., King, A. J., Conradt, L., Kerth, G., Lusseau, D., Mettke-Hofmann, C., Schaffner, C. M., Williams, L., Zinner, D., & Aureli, F. (2011). Collective decision-making and fission– fusion dynamics: A conceptual framework. Oikos, 120(11), 1608–1617.

Terborgh, J. (1990). Mixed flocks and polyspecific associations: Costs and benefits of mixed groups to birds and monkeys. American Journal of Primatology, 21(2), 87–100. 10.1002/ajp.1350210203

Thiollay, J.-M. (1999). Frequency of mixed species flocking in tropical forest birds and correlates of predation risk: An intertropical comparison. Journal of Avian Biology, 282–294.

Thrasher, D. J., Butcher, B. G., Campagna, L., Webster, M. S., & Lovette, I. J. (2018). Double-digest RAD sequencing outperforms microsatellite loci at assigning paternity and estimating relatedness: A proof of concept in a highly promiscuous bird. Molecular Ecology Resources, 18(5), 953–965. 10.1111/1755-0998.12771

Tóth, Z., Bókony, V., Lendvai, Á. Z., Szabó, K., Pénzes, Z., & Liker, A. (2009). Whom do the sparrows follow? The effect of kinship on social preference in house sparrow flocks. Behavioural Processes, 82(2), 173–177. 10.1016/j.beproc.2009.06.003

Wascher, C. A. F., Szipl, G., Boeckle, M., & Wilkinson, A. (2012). You sound familiar: Carrion crows can differentiate between the calls of known and unknown heterospecifics. Animal Cognition, 15(5), 1015–1019. 10.1007/s10071-012-0508-8

Whitehead, H. (2008). Analyzing animal societies: Quantitative methods for vertebrate social analysis. University of Chicago Press.

Wickham, H. (2011). Ggplot2. Wiley Interdisciplinary Reviews: Computational Statistics, 3(2), 180–185.

Wickham, H., Averick, M., Bryan, J., Chang, W., McGowan, L. D., François, R., Grolemund, G., Hayes, A., Henry, L., & Hester, J. (2019). Welcome to the Tidyverse. Journal of Open Source Software, 4(43), 1686.

Wrona, F. J. (1991). Group Size and Predation Risk: A Field Analysis of Encounter and Dilution Effects. The American Naturalist, 137(2), 186–201. 10.1086/285153

